# MRPS36 provides a missing link in the eukaryotic 2-oxoglutarate dehydrogenase complex for recruitment of E3 to the E2 core

**DOI:** 10.1101/2022.10.08.511390

**Authors:** Johannes F. Hevler, Pascal Albanese, Alfredo Cabrera-Orefice, Alisa Potter, Andris Jankevics, Jelena Misic, Richard A. Scheltema, Ulrich Brandt, Susanne Arnold, Albert J.R. Heck

## Abstract

The tricarboxylic acid (TCA) cycle, or Krebs cycle, is the central pathway of energy production in eukaryotic cells and plays a key part in aerobic respiration throughout all kingdoms of life. The enzymes involved in this cycle generate the reducing equivalents NADH and FADH2 by a series of enzymatic reactions, which are utilized by the electron transport chain to produce ATP. One of the pivotal enzymes in this cycle is 2-oxoglutarate dehydrogenase complex (OGDHC), which generates NADH by oxidative decarboxylation of 2-oxoglutarate to succinyl-CoA. OGDHC is a megadalton protein complex originally thought to be assembled just from three catalytically active subunits (E1o, E2o, E3). In fungi and animals, however, the protein MRPS36 has more recently been proposed as a putative additional component. Based on extensive XL-MS data obtained from measurements in mice and bovine heart mitochondria, supported by phylogenetic analyses, we provide evidence that MRPS36 is an essential member of OGDHC, albeit only in eukaryotes. Comparative sequence analysis and computational structure predictions reveal that in eukaryotic OGDHC, E2o does not contain the peripheral subunit-binding domain (PSBD), present in bacterial and archaeal E2o’s. We propose that in eukaryotes MRPS36 evolved as an E3 adaptor protein, functionally replacing the PSBD. We further provide a refined structural model of the complete eukaryotic OGDHC containing 16 E1o, 12 E3, and 6 subunits of MRPS36 accommodated around the OGDHC core composed of 24 E2o subunits (3.45 MDa). The model provides new insights into the OGDH complex topology and stipulates putative mechanistic implications.

## Introduction

TO maintain all necessary biological tasks of life, organisms and cells are in a constant demand for energy (1). To satisfy those needs, cells consume energy-rich fuels in a series of complex energy-transforming processes. At the center of aerobic energy metabolism are the enzymes of the tricarboxylic acid (TCA) cycle, or Krebs cycle, which in eukaryotes are located within mitochondria (2–4). The TCA cycle utilizes acetyl-CoA generated from sugars, fats and proteins in a series of enzymatic reactions to transfer electrons onto the reducing equivalents NADH and FADH2 (5). Subsequently, electrons are passed onto membrane bound respiratory chain protein complexes, fueling oxidative phosphorylation for the formation of adenosine triphosphate (ATP) (6, 7). Due to its central role in aerobic respiration, the TCA cycle has been extensively studied, resulting in a well-established view on enzymatic mechanisms and structural details of several of its enzymes (8–15).

Because of its sheer size and multi-component nature, the 2-oxoglutarate dehydrogenase complex (OGDHC) is a less well characterized key enzyme of the TCA cycle. OGDHC is a member of the 2-oxo acid dehydrogenase (OADH) family, alongside with the pyruvate dehydrogenase complex (PDC) and the branched-chain α-keto acid dehydrogenase complex (BCKDC). Each of these complexes consists of multiple copies of three catalytically active subunits (E1, E2, E3) assembling into multi-component enzymes weighing several megadaltons (MDa) that catalyze the oxidative decarboxylation of 2-oxo acids (16, 17). In contrast to E1 and E2, for which complex-specific genes exist (E1p/E1o/E1b, E2p/E2o/E2b), E3 is shared across all members of the OAHD complex family (18). Eukaryotic PDC exhibits an icosahedral core composed of 60 E2p subunits and 12 copies of an additional non-catalytic E3-binding protein (E3BP), OGDHC and BCKDC contain an octahedral core that is solely formed by E2o and E2b subunits, respectively (Figure 1). In all OADH complexes, E1 and E3 are thought to be arranged around the respective core, whereby the interaction involves a peripheral subunit-binding domain (PSBD) within E2. This relatively short (35-residues) PSBD is composed of two parallel alpha-helices (H1, H2) connected by an extended loop (19). Recruitment of E3 is mediated via charged side chains of H1 forming an electrostatic zipper (20).

**Fig. 1.**
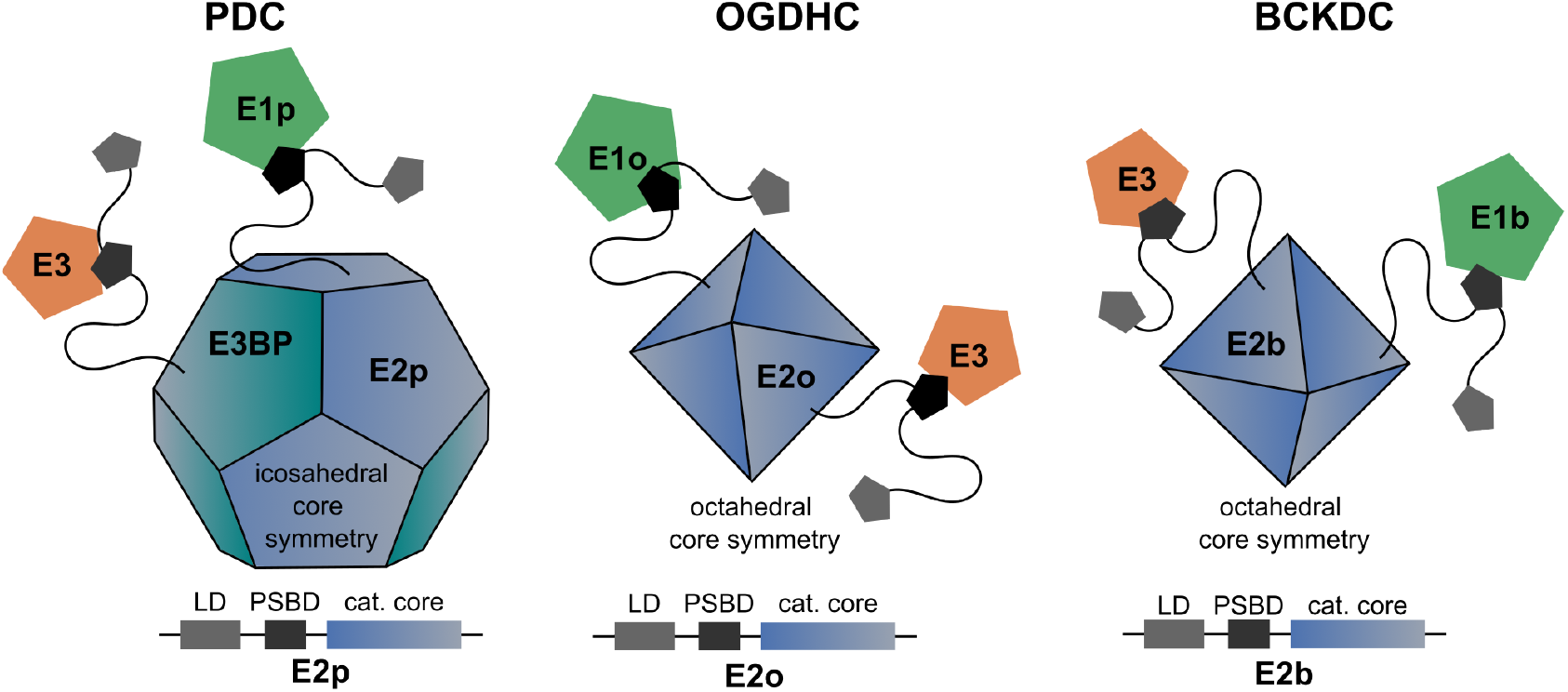
Schematic overview of the three members of the 2-oxo-acid dehydrogenase complex (OADHC) family. Pyruvate dehydrogenase complexes (PDC, left) contain an icosahedral E2p core to which E1p and E3 proteins become recruited. Additionally to E2p, PDC contains an alternative core forming subunit (E3BP) which specifically recruits E3. In contrast, for 2-oxoglutarate dehydrogenase complexes (OGDHC, middle) and branched-chain α-keto acid dehydrogenase complex (BCKDC, right) E1 and E3 are tethered around an octahedral E2 core. For all three OADH members, peripheral subunit-binding domains (PSBD) in the E2 sequence (bottom row) have been proposed to play a key role in recruiting E1 and E3 subunits to the core (21). For all family members, the lipoyl domain (LD) is essential for the carboxylation of 2-oxo acids, as it transfers respective intermediates from E1 to E2 components. Of these three, only OGDHC functions directly within the TCA cycle.

In contrast to PDC and BCKDC, OGDHC functions directly within the TCA cycle metabolizing 2-oxoglutarate to succinyl-CoA, CO2 and NADH + H+ in three consecutive steps. First, 2-oxoglutarate is decarboxylated by thiamine pyrophosphate (TPP)-containing E1o (OGDH) and the respective 2-succinyl intermediate is transferred to the flexible lipoyl domain (LD) of E2o (DLST). Second, E2o transfers the succinyl functional group from its LD domain onto CoA-SH generating succinyl-CoA. In a final reaction, the LD domain of E2o is re-oxidized by E3 (DLD), a FAD-containing subunit that transfers electrons to NAD+ producing NADH (22, 23). While this catalytic mechanism is well understood, current structural insights into OGDHC lack detailed information about the complex architecture (19, 24–26) in a close-to-native environment. Recent cryo-electron microscopic (cryo-EM) reconstructions of mammalian OGDHC revealed an octahedral core of 24 E2o subunits in an 8×3 assembly with E1o and E3 suggested to bind at the edges and faces, respectively (27, 28). Nonetheless, likely due to the highly conformational flexibility and possible heterogeneity of the complex (29), these structures are not sufficient to understand how the E1o and E3 components are organized with respect to the core. MRPS36 (KGD4), a small (11 kDa) and structurally unresolved protein, was more recently identified as a possible additional component of eukaryotic OGDHC (30–32). Its presence was suggested to be crucial for the efficient recruitment of E3 to the E2o core (32), but due to the lack of experimental evidence its function as an adaptor between E3 and the remainder of the OGDHC remains elusive. Also, in very recent high-resolution structural models of eukaryotic OGDHC, MRPS36 is not observed, or at least not reported on (27, 28).

Here, we explore the overall composition and architecture of mammalian OGDHC by combining complexome profiling (CP) (33, 34) and cross-linking mass-spectrometry (XL-MS) with heart mitochondria from both mice and bovine origin. Based on the XL-MS and CP data and supported by phylogenetic analyses, we propose that MRPS36 is an essential member of OGDHC, albeit exclusively in eukaryotic mitochondria. Comparative sequence analysis reveals that MRPS36 replaces a functional domain, termed PSBD, that is preserved across prokaryotic E2o proteins and the other OADH complex members (PDC, BCKDC), but is absent in eukaryotic E2o. Based on our data and computational modeling, we provide a refined structural model of the eukaryotic OGDHC highlighting how E1o, E3 and MRPS36 are organized with respect to the core of E2o subunits and how they assemble into a functional complex of over 3.45 MDa to act as a crucial component of the TCA cycle.

## Result and Discussion

### MRPS36 is a genuine member of OGDHC interacting with both the E2o core and E3

To delve into the architecture of eukaryotic mitochondrial OGDHC and to verify the presence of MRPS36, we examined XL-MS data obtained from intact bovine heart mitochondria (BHM) (35) and murine heart mitochondria (MHM) (**Supplementary Table 1**). The here obtained XL-MS data for OGDHC of MHM were extended with previously published XL-MS data from our laboratory (36). In both, mouse and bovine heart mitochondria, a substantial number of inter and intra-cross-links were observed for known members of the OGDHC (E1o, E2o, E3) and, notably, also for MRPS36. Our cross-linking experiments were more exhaustive for the BHM sample (as we applied three different chemical cross-linker reagents: DSSO, PhoX and DMTMM), resulting in a higher number of identified cross-links than for MHM (only cross-linked with DSSO). The observed cross-link patterns for all OGDHC subunits, including MRPS36, showed a high consistency between all technical and biological replicate datasets obtained from mitochondria of the two different organisms and by using different cross-linkers (**Figure 2A**, **Supplementary Table 1**). On a separate note, we did not observe any crosslinks between MRPS36 and mitochondrial ribosomal proteins, providing further evidence that its initial name as mitochondrial ribosomal protein S36, is indeed a misdemeanor. Several cross-links between all three components (E1o-E2o, E1o-E3, E2o-E3) were identified providing insights into the architecture of the binding interfaces within OGDHC. Our data showed that E1o interacts via its N-terminus and catalytic domain with both, a linker region (C-terminally of the LD-domain) and the catalytic domain of E2o. The involvement of respective regions in the formation of an E1o-E2o subcomplex as detected by XL-MS, is consistent with NMR and HDX-MS data reported for human OGDHC (38). Links between E1o and E3 components were also observed, albeit only in MHM, supporting previous findings that interactions between E1o and E3 are substantially weaker (38). Such weak E1o-E3 interactions could also hint at the absence of a stable subcomplex, which seems not surprising given that there is no direct dependency between the catalytic activities of E1o and E3. Our XL-MS data revealed stable interactions between residues of the LD-domain of E2o with the adjacent flexible linker region and the FAD/NAD binding domain of E3. In bacterial PDC and OGDHC, E3 is recruited to the E2 core via the PSBD of E2 (19–21, 39, 40). For eukaryotic OGDHC, such an E2-PSBD was postulated and proposed to be located in the flexible linker region, C-terminal to the LD-domain (27). However, so far, there has been no evidence found corroborating this assumption.

**Fig. 2.**
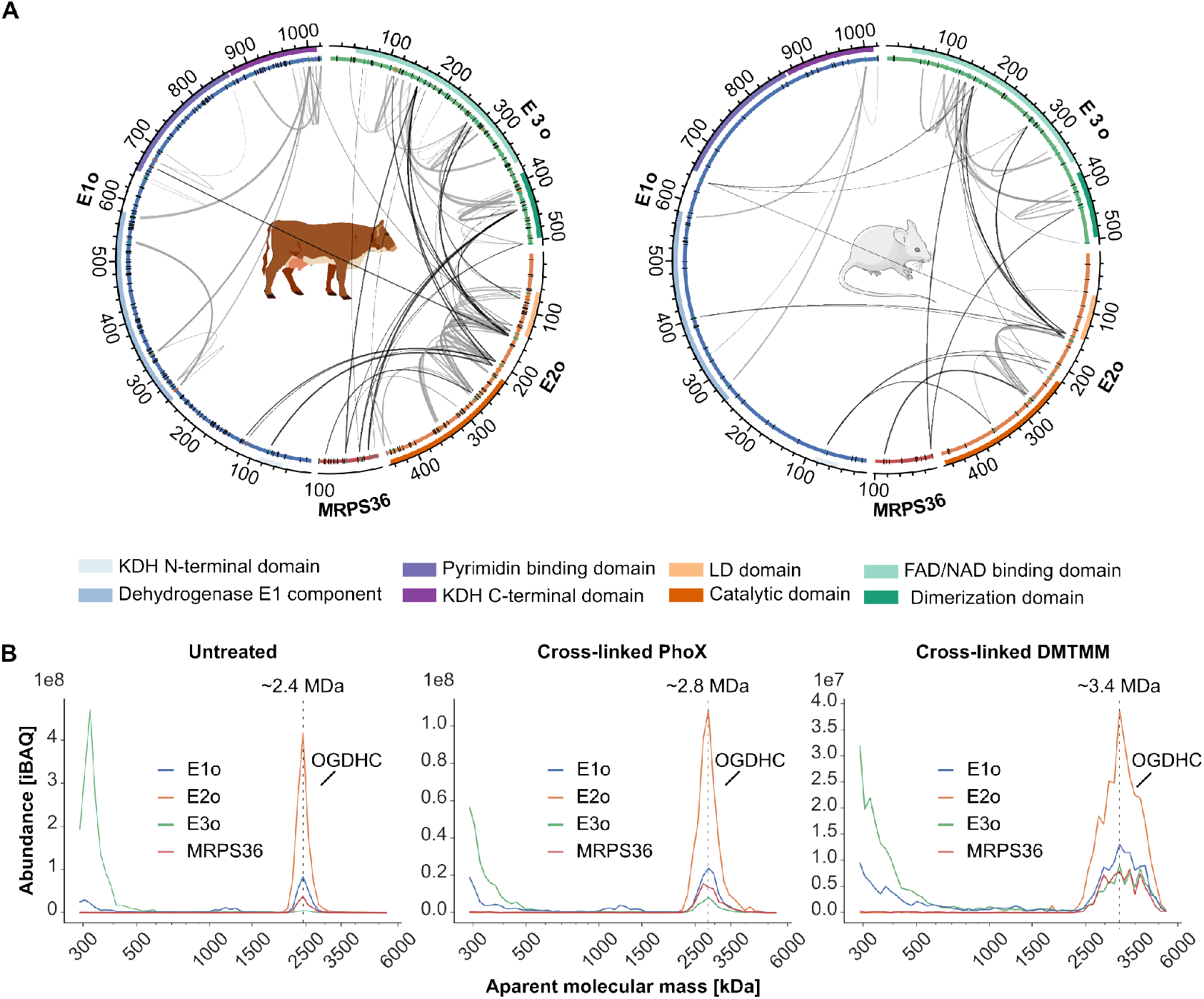
Exploring the architecture of eukaryotic OGDHC using XL-MS and CP. **A.** Circos plots showing intra and inter cross-links for members of OGDHC from bovine heart mitochondria (BHM, left) and mouse heart mitochondria (MHM, right). BHM were cross-linked with three different chemical cross-linkers (DSSO, PhoX and DMTMM) providing Lys-Lys and Lys-Asp/Glu cross-links. MHM were only cross-linked by using DSSO providing Lys-Lys cross-links. Cross-links for MHM were merged with previously published data from our lab (36). Functional domains are highlighted as colored tracks and were annotated using Interpro (37). Cross-linkable residues are indicated in a separate track. Depending on whether they are engaged in a cross-link they are either black (not cross-linked) or colored (involved in a cross-link) with Lys residues color-coded green and Asp and Glu orange. Sequences for E1o, E2o and E3 include mitochondrial transit peptides **B.** Migration profiles of bovine OGDHC subunits from non–cross-linked (untreated) and cross-linked (PhoX or DMTMM) heart mitochondrial samples separated by BN-PAGE (3 to 10%). MRPS36 is found to co-migrate with E1o, E2o, E3 subunits of the OGDHC. Peaks are annotated based on the apparent molecular masses of monomeric E1o (111 kDa), monomeric E2o (41 kDa), monomeric E3 (50 kDa) and monomeric MRPS36 (11 kDa). Cross-linking seems to stabilize binding of E3 to OGDHC increasing the apparent molecular mass of the complex. From the untreated samples it may be extracted that MRPS36 seems to bind to OGDHC even when E3 seems to be nearly absent

Most interestingly, in the XL-MS data from both samples (MHM and BHM), MRPS36 was found to be cross-linked to E2o and E3 by several inter-links, pointing to MRPS36 as a subunit of eukaryotic OGDHC. The N-terminal residues of MRPS36 were cross-linked to the FAD/NAD binding domain and dimerization domain of E3. In contrast, interactions to the E2o component involved C-terminal residues of MRPS36. In BHM, E2o and E3 share an interaction site on MRPS36 (K57). Although a direct interaction between those proteins has been hypothesized (32), no cross-links were identified between MRPS36 and E1o.

To corroborate our XL-MS findings, we also evaluated the architecture of the OGDHC by complexome profiling (CP) analyzing both naïve (untreated) as well as cross-linked (by either PhoX or DMTMM) BHM (**Supplementary Table 2**). In line with our XL-MS data, MRPS36 was found to comigrate with E1o, E2o, and E3 subunits of the OGDHC (**Figure 2B**). The four protein components were found to always co-migrate, albeit that the relative abundance of specific components varied substantially depending on the treatment (**SI Figure 1A**). Notably, we observed that the OGDHC eluted at different apparent molecular masses (untreated sample: 2.4 MDa, PhoX treated sample: 2.8 MDa, DMTMM treated sample: 3.4 MDa). These differences observed in apparent mass likely reflect the number of copies of E3 and MRPS36 subunits in the complex and possibly also to a lesser extent E1o (**Figure 2B**, **SI Figure 1A**). Substantial incorporation of E3 into the OGDHC was primarily observed in the cross-linked samples, further supporting the earlier notion that E3 interactions with E2o are weaker than E1o-E2o interactions (38). The apparent mass for respective OGDHC peaks in CP were increased after cross-linking thus suggests that more copies of E1o, E3 and MRPS36 subunits were stably incorporated into the complex upon chemical fixation (**SI Figure 1A**). The CP data suggest that E1o and MRPS36 are associated with the E2o core, even at low amounts of incorporated E3. Inspecting previously deposited CP-MS data retrieved from the ComplexomE profiling DAta Resource (CEDAR) repository (41) (Identifier: CRX34) and XL-MS data of human mitochondria (42) (**Supplementary Table 1**, **Supplementary Table 2**), we retrieved additional evidence confirming that MRPS36 is a genuine member of eukaryotic OGDHC (**SI Figure 1B-C**). The exact stoichiometry for the components of the OGDHC assembly is still unknown. Earlier reports on human OGDHC reported an approximate stoichiometry of 3(E1o)2:3(E2o)3:1(E3)2, but this report did not take the presence of MRPS36 into account (38). As crosslinking with DMTMM stabilized the OGDH complex best (**SI Figure 1A**), we estimate an apparent mass of 3.4 MDa for a fully assembled eukaryotic complex.

Taken together, XL-MS and CP data unambiguously demonstrate that MRPS36 is a key member of eukaryotic OGDHC directly engaging with E2o and E3 through its C-terminus and N-terminus, respectively. XL-MS provides unprecedented insight into so-far unresolved protein interactions within the eukaryotic OGDHC. Cross-linking the sample prior to CP-MS analysis preserves binding of E1o, E3 and MRPS36 to the E2o core suggesting an apparent mass of 3.4 MDa for the fully assembled BHM OGDHC.

### The evolutionary path of MRPS36 provides insights into its functional role

To dissect the functional role of MRPS36 as a fourth subunit of eukaryotic OGDHC, we investigated its evolutionary origin. Since E2o subunits form the core of all OGDH complexes, to which E1o and E3 are recruited (18), we included it into the phylogenetic analysis (**Figure 3A**). We observed that across the kingdoms of life, homologs of E2o, as well as E1o and E3, could be identified across all three branches of life, while homologs of MRPS36 are identified exclusively in eukaryotes (**Figure 3A**, **SI Figure 2A**). This finding suggests that MRPS36 is a protein exclusive to mitochondria (43). Notably, E2o displayed a much lower variability in eukaryotes than in bacteria and archaea. This reduced diversification suggests, that MRPS36 and E2o in mitochondria converged towards a tethered, and possibly reversibly controlled functional interaction (**Figure 3A**). It should be noted that the annotation for E2o is poor for archaea but assuming that they also harbor an OGDH complex, in addition to PDC and BCDCK (44), it was possible to tentatively assign entries to OGDHC based on similarity of the E2 catalytic domain to the bacterial and eukaryotic orthologues (**Figure 3B**, **SI Data**).

**Fig. 3.**
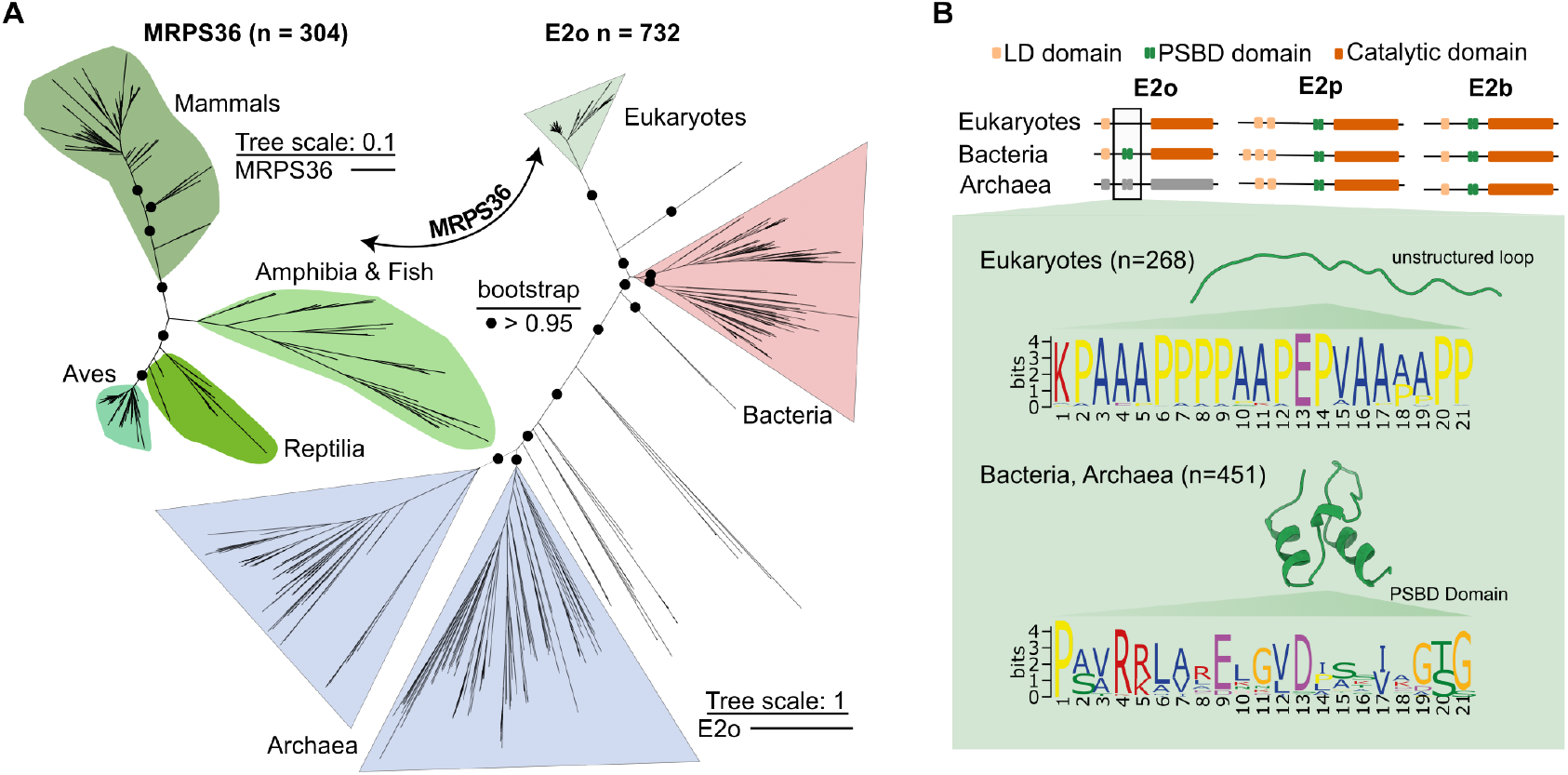
Phylogenetic and sequence analysis of MRPS36 and E2o. **A.** Unrooted maximum likelihood phylogeny trees are reported (IQ-TREE, substitution models LG+F+R10 for E2o and VT+F+R8 for MRPS36) based on 304 homologous sequences from eukaryotes for MRPS36 and 732 homologs of E2o selected from among eukaryotes, archaea and bacteria species. Node supports are indicated as circles if bootstrap support (200 replicates) is >95% only on the main branches (**SI Data**). The scale bar represents the average number of substitutions per site. **B.** Schematic view of the annotated domains for the 2-oxo-acid dehydrogenase (OADH) complex family in the three kingdoms of life, highlighting the presence of the PSBD sequence (green) in both E2p and E2b across all kingdoms of life, but the absence of PSBD in eukaryotic E2o proteins (missing green box) and the presence of such a domain in E2o from bacteria and likely also archaea. Notably, as shown E2p and E2b contain PSBD in all three kingdoms of life. E2o of archaea is colored gray, as relative sequence annotations are low, however clustered sequences share sequence features (including the catalytic domain) that clearly qualify them as E2o homologs. In bacterial and archaeal E2o proteins, a conserved sequence pattern is observed which corresponds to the structured PSBD as depicted in a recently published structural model (20) (PDB: 1EBD). In the eukaryotic E2o sequences, the motif in the corresponding stretch is clearly distinct with a conservation of proline and alanine residues that form an unstructured loop as predicted by AlphaFold2 (**SI Figure 2B**). The logos were generated using the MEME Suite (45)

A more detailed analysis at the sequence level revealed that the characteristic sequence pattern for PSBD is missing in eukaryotic E2o, and was replaced by an unstructured loop containing multiple alanine and proline residues (Figure 3B, SI Figure 2B, SI Data). Since this PSBD is crucial for the recruitment of E3 to the E2p core PDCs (20, 46–48), another mechanism mediating this functionally critical interaction has to be postulated for eukaryotic OGDHC.

It is therefore tempting to speculate that MRPS36 may have emerged as a small adaptor protein that specifically recruits E3 to the eukaryotic OGDHC.

It is worth noting that, in contrast to OGDHC, eukaryotic PDC adopted a different strategy to recruit E3 to its E2 core by developing a catalytically inactive paralogue E2 called E3BP that contains a PSBD (49, 50). These specialized mechanisms to recruit E3 may be important to adapt its distribution to different metabolic requirements. As such, it would be interesting to investigate further whether such a strategy also exist for eukaryotic BCKDC, the third member of the 2-oxo acid dehydrogenase (OADH) complex family.

In summary, based on our phylogenetic analysis we propose that MRPS36 evolved to substitute the missing E2-PSBD of OGDHC that is absent in eukaryotic E2o. The presence of MRPS36 may enable specific recruitment of E3 to E2o in a controllable fashion.

### MRPS36 is essential for the recruitment of E3 to E2o in eukaryotic OGDHC

To corroborate our hypothesis that the loss of PSBD enforced changes in the eukaryotic E2o to E3 binding, we set out to structurally characterize the E2o-E3 interactions using the structural prediction algorithm AlphaFold2-Multimer (51) (AF2). First, we constructed an interface between E2o and E3 for *B. taurus* (an E2o not containing a PSBD) and *E. coli* (an E2o harboring a PSBD) (**Figure 4A**). For both systems, the AlphaFold2 predictions confidently recapitulated known protein domains for E2o and E3 (**see also Figure 3B**) (28, 52). In all top five models, the lipoylated lysine (K43) of the LD-domain was located in close proximity to the respective catalytic domain (**Figure 4A**, **SI Data**). A similar orientation of the LD-domain was observed recently for the E2p-LD domain of the bacterial PDC (53). Interestingly, only the *E. coli* E2o-E3 complex – and not the eukaryotic complex - formed an interface, with the involved E2o residues being predominately located within the H1 helix of the PSBD, as previously reported for the bacterial E2 core (E2p) of PDC (20). The absence of a predicted E2o-E3 binding interface in *B. taurus* supports our hypothesis that in eukaryotes E3 binding to E2o is hampered and may thus rely on an additional adaptor protein. Therefore, we set out to structurally characterize and model MRPS36 as a possible additional subunit of eukaryotic OGDHC to facilitate the binding of functional assemblies of E3 (dimeric) to E2o (trimeric) (27, 28, 54). The final structural model highlights how dimeric E3 may be recruited to the trimeric E2o core via MRPS36 (**Figure 4B**). In our model, MRPS36 interacts via its C- and N-terminal residues with the two E2o subunits of the E2o trimeric core and with both E3 subunits of the E3 dimer, respectively. In eukaryotic OGDHC, MRPS36 occupies a significant part of the binding interface that was reported earlier for the interaction of E2p-PSBD-with E3 in bacteria (**SI Figure 3A**) (20). This supports the hypothesis that MRPS36 evolved as an additional OGDHC subunit, functionally replacing the missing PSBD in eukaryotic E2o.

**Fig. 4.**
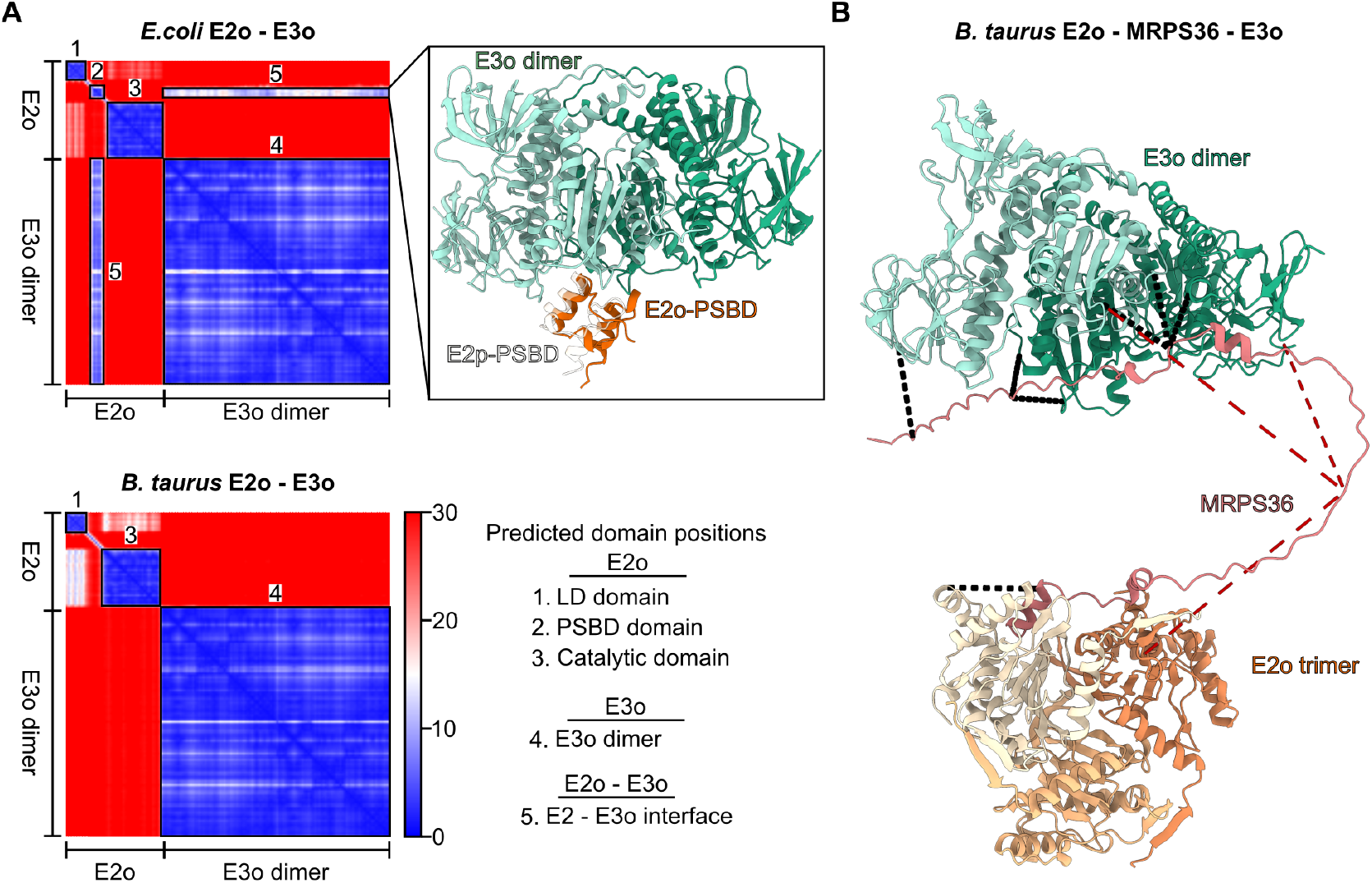
In eukaryotic OGDHC, MRPS36 is essential for the recruitment of E3 to E2o. **A.** The predicted aligned error (PAE) plots of AlphaFold2 predictions of the E2o-E3 interaction within eukaryotic (*B. taurus*) and prokaryotic (*E. coli*) model systems. The PAE plot describes the confidence in the relative positions and orientations for residues (in Angstrom) in the prediction, thereby describing intra and inter domain orientations. The smallest possible value of the PAE is 0 Å. For both predictions, AlphaFold2 confidently predicts relative residue orientations for known domains with high confidence as indicated by the black, numbered boxes. Domain and interface annotations for both AlphaFold2 predictions are listed as “Predicted domain positions”. For *E. coli* E2o-E3 AlphaFold2 predicts two additional high confidence regions corresponding to a PSBD (box 3) and an inter domain interface corresponding to E2-PSBD and the respective E3 subunits. The structural details of the interaction are further highlighted in box 5. Interaction partners are colored as indicated. To further validate the predicted interaction, the previously published crystal structure for the E2p-PSBD and E3 complex of a bacterial PDC(Mande et al., 1996) (PDB: 1EBD) is structurally aligned to E3 as reference. **B.** AlphaFold2 prediction highlighting structural details of how MRPS36 recruits E3 to E2o in a functional edge (E2o trimer; E3 dimer) of a eukaryotic OGDH complex. In the model, E3 subunits are colored in different shades of green, E2o subunits in different shades of orange and MRPS36 is colored red. Obtained cross-links (see Figure 1A) are mapped onto the final complex with cross-links <30 Å being colored black and cross-links >30 Å being colored red. For clarity, only residues corresponding to the catalytic domain of E2o are shown (residues 155-387)

Next, we investigated the sequence conservation of MRPS36 (**SI Figure 3B**). Notably, conserved residues of MRPS36 are observed at the interfaces to E2o and E3 suggesting the functional importance of these residues and further supporting the model. The binding of MRPS36 to E2o and E3 seems predominately stabilized by electrostatic interactions (SI Figure 3C), similarly to the binding of E2p-PSBD to E3 (20). Additionally, the final structural model agrees well with the XL-MS data (**Figure 4B**, **Supplementary Table 1**), except for those cross-links involving MRPS36-K58, the interaction site shared between E2o and E3, that do not fulfill the threshold distance constraint of 30 Å. This is of interest as this site resides close to a serine residue (S61) that we found phosphorylated in BHM (**SI Figure 3D**). While S61 is not part of the highly conserved N- and C-termini, further sequence analysis revealed a prominent motif in vertebrates, with serine residues being an integral part of the sequence stretch that is connecting the termini of MRPS36 (**SI Figure 3D**).

In line with this observation, the serine residue S61 has also been found to be phosphorylated in the corresponding human (S61) and mouse (S60) residues in recent phosphoproteomic studies (55–57). Further studies are needed to elucidate whether this phosphorylation site impacts the interaction of OGDHC subunits and thus its regulation.

In summary, our computational modeling supports the conclusions drawn from our phylogenetic analyses, highlighting the importance of MRPS36 as a functional substitute for the PSBD that is absent in eukaryotic E2o. The structural model agrees well with the experimentally generated XL-MS restraints and reveals structural details of how MRPS36 mediates binding of E3 to the eukaryotic OGDHC core.

### Towards a refined complete structural model of eukaryotic OGDHC

Several recently published structural models of eukaryotic OGDHC revealed an octahedral core containing 24 E2o subunits (27–29). However, these models do not provide comprehensive details about the exact orientation of E1o and E3 with respect to the core, and fully overlooked the presence and role of MRPS36 (27–29). To understand how all subunits of the eukaryotic OGDHC are assembled into a complex, we set out to use AlphaFold2-Multimer (51) together with our XL-MS data to model the orientations of E1o, E3, and MRPS36 with respect to the E2o core, and to predict potential interfaces. To do so, we first modeled the so far unresolved interactions between E1o and E2o, using the previously described functional assembly of an E2o trimer and an E1o dimer as a starting point (21, 58, 59). In the resulting model, fully supported by XL-MS data (**see Figure 2A**, **Supplementary Table 1**), the E1o dimer is tethered to the E2o core via its N-terminus and is interacting with the LD domains of E2o subunits.

To build a structural model that reflects all protein-protein interactions of OGDHC we further modelled the LD(E2o)-E3 interaction, thereby providing structural insights of final catalytic reaction of OGDHC - the regeneration of the disulfide bridge in the lipoyl-group in the LD domain of E2o. Finally, we used these generated structural interfaces to assemble a complete eukaryotic OGDHC model by structurally aligning E2o subunits to a previously published cryo-electron microscopic structure of the human E2o core (28) (**Figure 5A**). In the final model, 16 E1o subunits, 12 E3 subunits, and 6 subunits of MRPS36 are accommodated around the OGDHC core composed of 24 E2o subunits (**Figure 5B**). The calculated molecular mass for this model of eukaryotic OGDHC is 3.45 MDa and thus in good agreement with the apparent massed observed in our CP-MS experiments when the binding of components E1o, E3 and MRPS36 was stabilized by XL using DMTMM (**Figure 2B**). Further, our XL-MS data are in very good agreement with the assembled OGDHC model, with the median distance for mapped cross-links being shorter than 20 Å (**SI Figure 4A**, **Supplementary Table 1**). Cross-links exceeding the cut-off distance of 30 Å involve mostly highly flexible linker regions of E2o connecting the LD-domain to the catalytic domain (**SI Figure 4A**).

**Fig. 5.**
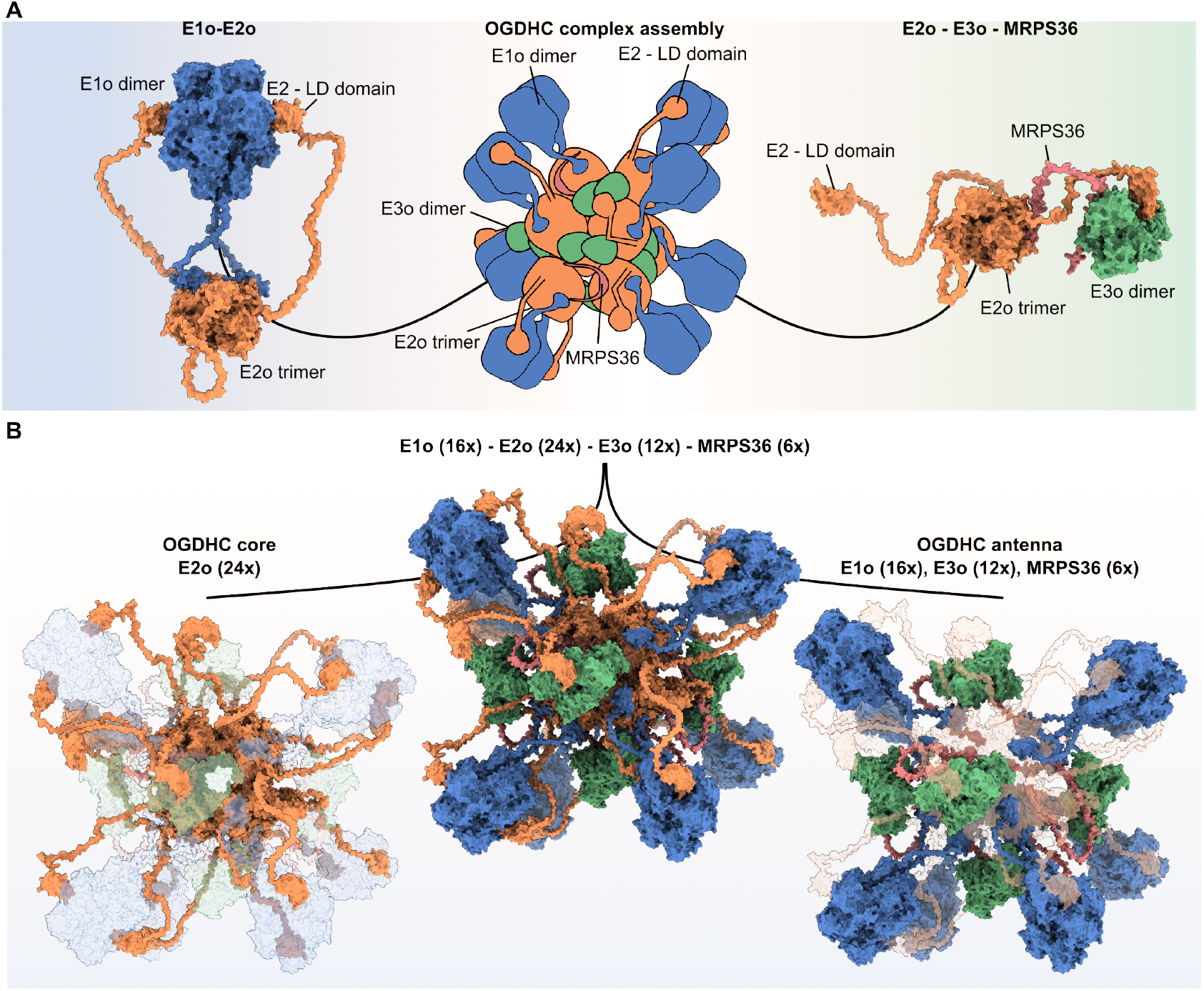
Assembly of an all-inclusive structural model of eukaryotic OGDHC. **A.** Modelled complex interactions for E1o (dimer, blue) - E2o (trimer, orange) (left) and E2o (trimer, orange) – E3 (dimer, green) – MRPS36 (single chain, red) (right). Structural models were built on the basis of a cryo-electron microscopy structure of the human E2o core (28) (**PDB: 6H05**) as indicated in the overview (center). **B.** All-inclusive model of the eukaryotic OGDHC (*B. taurus*) (center). The OGDHC core is composed of 24 E2o subunits assembled into 8 trimers (orange, left panel). The OGDHC antenna is composed of 16 E1osubunits assembled into 8 dimers (blue), 12 E3 subunits assembled into 6 dimers (green) and 6 x MRPS36 (red) connecting each E3 dimer to a E2o trimer

The assembled OGDHC model provides novel insights into the interaction of the LD(E2o) domains with the catalytic domain of E1o and E3 (**SI Figure 4B**). For both interfaces, the lipoyl-group of the LD domain (modeled as LA2 on Lys43) is at an appropriate distance to its respective co-factors in E1o (TPP, 8 Å) and E3 (FAD+, 15 Å). Furthermore, for the E2o-LD-E1o interface, residues His750 and His473 of the E1o subunits are sufficiently close to the lipoyl group (3 Å and 5 Å, respectively) to act as proton acceptor after nucleophilic attack of one sulfur atom resulting in the transfer of decarboxylated succinyl intermediate to E2o via this lipoyl group (18, 60). Detecting transient interactions, such as those involving the E2o-LD domain, is generally thought to be extremely challenging to predict using computational approaches (61). However, we were able to identify all observed protein-protein interactions when refining predictions based on our experimental constraints from the XL-MS analysis (**see Material and Methods for more information**). Further, the here shown data highlight how XL-MS and CP-MS in combination with computational modeling can help to overcome positional ambiguity in modest-resolution cryo-EM density maps, as observed for a cryo-EM model of bovine OGDHC reported recently(27).

## Conclusions

Based on extensive cross-linking data obtained from both mouse and bovine heart mitochondria, and further supported by complexome profiling data, we identify MRPS36 as a key subunit of OGDHC, exclusively present in eukaryotes. We provide evidence that MRPS36 evolved as an E3 adaptor protein, functionally replacing the PSBD of prokaryotic E2o. Consequently, MRPS36 mediates the interaction between E2o and E3. Using integrative structural approaches, we provide a refined structural model detailing on how MRPS36 mediates binding of E3 to the eukaryotic OGDHC E2o core. We expanded the model by including all intra- and interprotein interactions detected by XL-MS, ultimately producing a compelling eukaryotic OGDH complex of 3.45 MDa that contains 24 E2o subunits, 16 E1o subunits, 12 E3 subunits, and 6 MRPS36 subunits. Moreover, as XL-MS can be performed in naïve intact mitochondria and as shown here, can stabilize weaker interactions it uniquely complements structural efforts that require protein samples to be transferred into detergents, which may affect the structurally stability and topology of certain complexes. The complete structural model of eukaryotic OGDHC presented here can be used to extract so far unrevealed structural insights into the enzymatic reactions by each of the catalytically active sites. Overall, this sheds further light on mechanistic details of the eukaryotic mitochondrial OGDHC and its importance within the TCA cycle.

## ACKNOWLEDGEMENTS

All authors acknowledge support from the Netherlands Organization for Scientific Research (NWO) funding the Netherlands Proteomics Centre through the X-omics Road Map program (Project 184.034.019) and the EU Horizon 2020 program Epic-XS (Project 823839). UB and SA were supported by the Netherlands Organization for Health Research and Development (ZonMW Project 91217009) and the German Research Foundation (DFG) through the Collaborative Research Center 1218 (Project 269925409). JFH and PA acknowledge the Dutch National Supercomputer, supported by NWO, for the computational resources (grant agreement EINF-894). We thank Dusanka Milenkovic and Nils-Göran Larsson for their expertise and assistance throughout all aspects of our study and for reviewing the manuscript.

## AUTHOR CONTRIBUTIONS

JFH and AJRH conceptualized the study. JFH performed the XL-MS experiments with support from JM. JFH performed the XL-MS analysis and structural modeling with support from PA, AJ, and RAS. PA performed phylogenetic- and sequence analysis. ACO and AP performed the complexome profiling experiments for the bovine and human mitochondrial samples. ACO, SA and UB analyzed the complexome profiling datasets. ACO and SA provided the bovine mitochondrial samples. JM provided the mouse heart mitochondrial samples. JFH, and AJRH wrote the original draft, with all authors editing the manuscript before submission. UB, SA, and RAS. AJRH acquired funding and resources. AJRH supervised the project.

## COMPETING FINANCIAL INTERESTS

The authors do not declare any conflict of interest.

## Material and Methods

### Bovine heart mitochondria (BHM)

All experimental XL-MS and CP-MS data for OGDHC from bovine heart mitochondria presented in this manuscript were previously generated and are published by us. Detailed information about the sample preparation, cross-linking as well as complexome profiling can be found in the published manuscript entitled “*Molecular characterization of a complex of apoptosis-inducing factor 1 with cytochrome c oxidase of the mitochondrial respiratory chain”* (35). Raw data for XL-MS and CP-MS are publicly available at the ProteomeXchange partner PRoteomics IDEntifications (PRIDE) database and the ComplexomE profiling DAta Resource (CEDAR) database with the identifiers PXD025102 and CRX33, respectively.

### Identification of MRPS36 phosphorylation sites in BHM

24 SCX fractions corresponding to BHM peptides cross-linked with DSSO (35) were analyzed using a classical bottom-up workflow. Briefly, fractions were injected in an Agilent 1290 Infinity UHPLC system (Agilent) on a 50cm analytical column packed with C18 beads (Dr Maisch Reprosil C18, 3 μm) coupled online to a Q Executive HF (Thermo Fisher Scientific). We used the following LC-MS/MS parameters: after 5 minutes of loading with 100% buffer A (water with 0.1% formic acid), peptides were eluted at 300 nL/min with a 80 minutes gradient from 4% to 39% of buffer B (80% Acetonitrile and 20% water with 0.1% formic acid). For MS acquisition we used a MS1 Orbitrap scan at 120,000 resolution from 300 to 1600, AGC target of 3e6 ions and maximum injection time of 120 ms. The ions with a charge from +2 to +8 were fragmented (NCE of 27%) and analyzed with MS2 Orbitrap at 30,000 resolution, AGC target of 1e5 ions and maximum injection time of 75 ms. Respective spectra were afterwards analyses with MQ (62) using following settings: Enzyme: (Trypsin), Oxidation (M); Acetyl (Protein N-term); Phosphorylation (STY); Carbamidomethylation (C). Identified peptides as well as phosphorylation sites of MRPS36 were next visualized using Alphamap (63). A sequence motif analysis for residues around the reported phosphorylation site (S61) was performed for homologue sequences for MRPS36 from vertebrates (SI Data) using MEME Suite (45).

### Structural modeling of E2 proteins of *Bos taurus*

Structural models of E2 proteins of the PDC, OGDHC and BCKDC were generated using AlphaFold2 (version 2.2.0, with model preset set as monomer_ptm) (64). For each prediction five models were generated. Predicted aligned error (PAE) plot and a per-residue estimate of its confidence on a scale from 0 - 100 (pLDDT) plot were using a custom python script.

### Structural modeling of OGDHC component interactions and building of a complete OGDHC model

Protein-protein interactions were modelled with AlphaFold2-Multimer (51) (version 2.2.0, template database date: 2021-11-01). For each prediction 25 models were generated. Predicted aligned error (PAE) plot and a per-residue estimate of its confidence on a scale from 0 - 100 (pLDDT) plot were generated for the top 5 ranked models using a custom python script. Predictions of protein-protein interactions were performed with previously reported functional assemblies of respective eukaryotic OGDHC components (E1o dimer, E2o trimer, E3 dimer) (28, 54, 58, 59). A final complex for E2o-E3-MRPS36 was assembled from models generated for E2o-MRPS36 and E3-MRPS36. Briefly, respective models were aligned on MRPS36 and the flexible loop was refined using Modeller (65) in Chimera 1.14 (66). Modelling the LD(E2o) interaction with the catalytic domain of E3 dimer, was performed with sequences that correspond to the LD (E2o) and dimeric E3, including all residues that were found to be crosslinked. Similarly, the full-length model, connecting the LD-domain to the catalytic domain of E2o was build. To map distances of the lipoyl residue of the LD domain (K43), a LD-domain containing a lipoyllysine (LA2) was modeled similar as previously described (67). Briefly, LA2 was added to the previously modelled LD-domain by superimposing the LA2 from the PDB ligand library onto the K43 and connected using ChimeraX (68). Torsion angles were refined in Coot. A final OGDHC model was built in ChimeraX, by aligning previously produced models (E1o-E2o, E2o-E3-MRPS36) onto the previously published cryo-EM structure of a human E2o core (24 E2o subunits) (28) (PDB 6H05). A final complex model contains 16 Eo1 subunits, 24 E2o subunits, 12 E3 subunits and 6 MRPS36 subunits to a total mass of 3.4 MDa. Obtained clashes between the flexible linkers of E2o (connecting the LD-domain to the catalytic core) with other subunits of E1o, E2o and E3 were removed by modelling an alternative position using “Modell Loops” in ChimeraX (68, 69). Final visualization of modeled protein interactions, the final complex as well as mapping obtained cross-links onto respective structures was performed in ChimeraX. For mapping and distance analysis of the cross-links the XMAS tool (70) was utilized.

### Phylogenetic tree- and sequence motif analysis for E2o and MRPS36

Homologous protein sequences of E2o and MRPS36 of *B. taurus* were retrieved through NCBI-BLAST against the non-redundant protein sequences database (10-12 June 2022). For either proteins, three BLAST searches on 1000 targets were restricted to Eukaryotes, Archaea and Bacteria. At most four species per genus with a query coverage > 50% and a sequence identity < 90% were selected among the top 500 hits. For E2o, the resulting set of about 300 sequences for each kingdom was aligned with Clustal Omega (71) over 10 iterations and then manually curated using Gblocks (72) (SI Data). Maximum likelihood (ML) tree inference on the curated alignment set containing 732 distinct alignment patterns 63 invariant sites (7.7% of the total) was done with IQ-TREE (73) embedded in the galaxy webserver platform (74) with an optimized substitution model (LG+F+R10) over 100 bootstrap replicates. For MRPS36 the BLAST against Archaea and Bacteria did not produce any significant hit, therefore, 304 homologous sequences from Eukaryotic homologs were selected using the criteria described above for E2o to investigate the evolutionary relationships between the latter group. Maximum likelihood (ML) tree inference was conducted in the same fashion on polished alignment containing 128 amino-acid sites and 5 invariant sites (3.9% of the total) with an optimized substitution model (JTT+R5). Sequence motif analysis for the PSBD of E2o was done with the MEME Suite (45) using the aligned sequences corresponding to the region of the E2o PSBD of *E. coli*.

### Interface residue analysis MRPS36, E2o, E3

Residues of MRPS36 interacting with either E2o or E3 were assesses with Prodigy (75) using the E2o-E3-MRPS36 complex as input structure. A 2-D projection of the electro static surface potential of MRPS36 was generated using SURFMAP (76) using the predicted structural model of MRPS36 (pdb format) as well as a list of interacting residues as input. For E2o and E3 interacting residues, the electrostatic potential of the surface was calculated using the E2o-E3-MRPS36 complex in ChimeraX.

### Cross-linking of mouse heart mitochondria (MHM)

Mouse heart mitochondria purified from three biological replicates were diluted in MIB buffer to a protein concentration of 1 mg/mL and subsequently cross-linked with DSSO (Thermo Fischer Scientific) using optimized conditions (0.5 mM DSSO, 45 min at 15 °C). The cross-link reaction was quenched for 30 min at 25 °C by the addition of 50 mM Tris (1 M Tris buffer, pH 8.5). Cross-linked mitochondria were pelleted at 11,000 g at 4 °C for 10 min and re-suspended in mitochondrial lysis buffer (100 mM Tris pH 8.5, 7 M Urea, 1% Triton-X-100, 5 mM TCEP, 30 mM CAA, 2.5 mM Mg2+, proteinase inhibitor cocktail). Mitochondria were solubilized for 30 min on ice. Next, proteins in the soluble fraction were precipitated using Methanol/Chloroform precipitation as previously described (77). The dried protein pellet was then re-suspended in digestion buffer (100 mM Tris-HCl pH 8.5, 1% sodium deoxycholate, 5 mM TCEP, and 30 mM CAA). Protein digestion was performed overnight at 37°C using Trypsin (1:25 ratio weight/weight) and Lys-C (1:100 ratio weight/weight), respectively. Finally, peptides were desalted by solid-phase extraction C18 columns (Sep-Pak, Waters) and fractionated into 22 fractions using an Agilent 1200 HPLC pump system (Agilent) coupled to a strong cation exchange (SCX) separation column (Luna SCX 5 μm to 100 Å particles, 50 × 2 mm, Phenomenex). The 22 SCX fractions of DSSO were analyzed using an Ultimate3000 (Thermo Fisher Scientific) connected to a 50-cm analytical column packed with C18 beads (Dr Maisch Reprosil C18, 3 μm) heated at 45°C, connected to an Orbitrap Fusion Lumos. Peptides were eluted at 300 nL/min with a 95 minutes gradient from 9% to 40% of buffer B (80% Acetonitrile and 20% water with 0.1% formic acid). To identify cross-linked peptides, MS1 scans with the Orbitrap resolution set to 120,000 from 350 to 1400 m/z were performed (normalized AGC target set to 250% and maximum injection time set to auto). Ions with a charge from +3 to +8 were fragmented with stepped HCD (21%, 27%, 33%) and further analyzed (MS2) in the Orbitrap (30,000 resolution, normalized AGC target set to 200% and maximum injection time set to auto) for detection of DSSO signature peaks (difference in mass of 37.972 Da). The four ions with this specific difference were further sequenced following MS3 CID - Ion Trap scans (collision energy 35%, normalized AGC target set to 200% and maximum injection time set to auto). The raw files corresponding to the 22 DSSO fractions were analyzed with Proteome Discoverer software suite version 2.5 (Thermo Fisher Scientific) with the incorporated XlinkX node for analysis of cross-linked peptides as reported by Klykov et al. (78). Data were searched against a FASTA file containing proteins, which were previously identified following a classical bottom-up workflow. Where applicable, mitochondrial target peptides were removed from respective protein sequences. For the XlinkX search, fully tryptic digestion with three maximum missed cleavages, 10 ppm error for MS1, 20 ppm for MS2 and 0.5 Da for MS3 in Ion Trap was set as search parameters. Carbamidomethyl (C) was set as static modification while Oxidation (Met) and Acetylation (protein N-terminus) were set as dynamic modification. Cross-linked peptides were accepted with a minimum XlinkX score of 40 and maximum FDR (controlled at PSM level for cross-linked spectrum matches) rate set to 5%. Presented XL data for mouse OGDHC were supplemented with recently published data from our lab (36).

### Complexome profiling of human embryonic kidney (HEK) 293 mitochondria

HEK293 cells (ATCC CRL-1573) were cultured in DMEM (Lonza BE12–604F) supplemented with 10% fetal calf serum (GE Healthcare) in a 37°C incubator at 5% CO2 for 48 hours before the experiment. Cells were harvested by suspension in the same medium, pelleted and washed with ice-cold PBS. After recovery by centrifugation, cells were suspended in homogenization buffer (250 mM sucrose, 1 mM EDTA, 20 mM Tris-HCl, pH 7.4 supplemented with protease inhibitor cocktail (SIGMAFAST™)) and disrupted with 15 strokes using a Potter-Elvehjem homogenizer. Mitochondria were further isolated as described in (79). Crude mitochondria were purified using a two-layers sucrose gradient (1.0/1.5 M sucrose in 20 mM Tris-HCl pH 7.4, 1 mM EDTA) for 20 min centrifugation at 60,000 x g. After recovery from the interphase, pure mitochondria were washed in homogenization buffer and recovered by centrifugation (10,000 x g; 10 min; 4 °C). Protein concentration was determined by Bradford assay. Mitochondria (200 μg protein) were solubilized with digitonin (8g/g protein; SERVA) in 50 mM NaCl, 2 mM aminohexanoic acid, 1 mM EDTA, 50 mM imidazole, pH 7.0, and kept on ice for one hour. The lysate was cleared by centrifugation at 22,000 x g for 20 min; 4 °C. The supernatant was recovered and supplemented with loading buffer and separated on a 3-16% polyacrylamide gel by blue-native (BN)-PAGE as previously described (80). After the run, the gel was processed as described previously (81), except that the lane was cut in 56 even slices. In-gel trypsin digestion, peptide recovery, LC-MS and complexome profiling analysis were performed as described in (81) with slight modifications. Here, MS spectra were matched against the human reference proteome retrieved from UniProt using MaxQuant v.2.0.3; the mass calibration was done using human OXPHOS complexes and other well-known soluble proteins; data were analyzed using Microsoft Excel and R Studio.

## Data availability

Data for BHM XL-MS and CP-MS can be found at the ProteomeXchange partner PRoteomics IDEntifications (PRIDE) database with the identifier PXD025102 and the ComplexomE profiling DAta Resource (CEDAR) database with the identifier CRX33. CP-MS data for human HEK 293 mitochondria are available via CEDAR database with the identifier CRX34. The mass spectrometry proteomics data for BHM have been deposited to the ProteomeXchange Consortium via the PRIDE partner repository with the dataset identifier PXD036896. XL-MS data for MHM are available via the PRIDE partner repository with the dataset identifier PXD031345. AlphaFold2 models, sequences used for phylogenetic tree analysis and motif generation (SI Data) can be accessed via figshare.

## Notes

https://www.ebi.ac.uk/pride/

### Competing Interest Statement

The authors have declared no competing interest.

